# Changes in the Transcriptome and Synthetic Lethal Dependencies Following *KRAS* Mutant Expression Reveal Profound Tissue-Specificity

**DOI:** 10.64898/2026.04.30.721990

**Authors:** Timothy D Martin, Mei Yuk Choi, Julia C McBride, Stephen J Elledge

## Abstract

Oncogenic KRAS mutations exhibit a striking tissue-restricted tropism, occurring with high frequency in pancreatic, colorectal, and lung adenocarcinomas while remaining rare in other lineages. The molecular basis for why these specific tissues are uniquely permissive to KRAS transformation, and how this context shapes therapeutic vulnerabilities, remains poorly defined. Here, we utilized CRISPR-mediated genome engineering to generate endogenous, conditional KRAS-mutant isogenic cell line models across three primary permissive lineages (lung, colon, and pancreas) and the non-permissive breast lineage. Integrated genome-wide CRISPR fitness screens and comparative transcriptome analyses revealed that KRAS-driven synthetic lethal (SL) dependencies are profoundly shaped by their tissue of origin. Strikingly, we observed minimal overlap in SL hits across lineages, with only three genes shared among the permissive lines, suggesting that the KRAS oncogene operates through divergent, context-specific genetic networks. Mechanistically, we show that KRAS activation induces a universal MYC-driven metabolic signature, but the specific machinery required to sustain this state is lineage-restricted. We identified a dependency on the diphthamide synthesis pathway to maintain translational fidelity amidst a KRAS-induced hyper-translational state. These findings demonstrate that even when driven by the same oncogene, tumors exhibit distinct regulatory landscapes and unique genetic vulnerabilities. Our results provide a framework for developing lineage-aware therapeutic strategies, moving beyond universal KRAS inhibition toward targeted interventions tailored to a tumor’s specific tissue context.

**SIGNIFICANCE STATEMENT:** While *KRAS* mutations drive a significant portion of human malignancies, their prevalence is strikingly restricted to specific lineages, namely pancreatic, colorectal, and lung tissues. This tissue-restricted tropism suggests that oncogenic *KRAS* does not operate in a vacuum but requires a permissive, tissue-specific molecular landscape to sustain tumorigenesis. By integrating comparative transcriptome analyses with functional genomics across four isogenic lineages, we demonstrate that *KRAS* synthetic lethal dependencies are not universal but are hardwired to the cell of origin. This work establishes a framework for tissue lineage-aware oncology, shifting treatment paradigms from targeting the *KRAS* mutation alone to targeting the specific genetic networks, defined by the tissue of origin, that sustain KRAS-driven growth.

## INTRODUCTION

The RAS gene family (*HRAS, KRAS*, and *NRAS*) comprises the most frequently mutated oncogenes in human cancer. These mutations typically impair intrinsic GTPase activity and confer resistance to GTPase Activating Protein (GAP) mediated hydrolysis, locking the RAS protein in a constitutively GTP-bound, active state that drives aberrant cell proliferation and survival^1^.

Historically, the structural lack of “druggable” pockets on the KRAS surface earned it the reputation of being “undruggable”^2^ Recent breakthroughs in covalent chemistry have led to the FDA approval of direct inhibitors; however, clinical efficacy remains limited by rapid resistance in many tumor types^3,4^. More recently, non-covalent pan-KRAS inhibitors like RMC-6236/Daraxonrasib have been developed that work by sterically preventing downstream KRAS signaling due formation of a tri-complex comprised of the inhibitor with GTP-bound KRAS with cyclophilin A. These inhibitors are not dependent on a particular codon mutation and are already showing clinical promise^5^. An alternative therapeutic strategy involves identifying KRAS synthetic lethal (SL) partners. These SL partners represent genetic dependencies that are essential only in the presence of oncogenic KRAS, offering a therapeutic window to selectively eliminate mutant cells while minimizing toxicity to wild-type (WT) tissues.

While previous RNAi^6^ and CRISPR-based^7^ screens have identified several KRAS dependencies, including mitotic^8^ and mitochondrial^9^ pathways, many other hits often fail to translate across different clinical contexts. *KRAS* mutations are remarkably restricted to the pancreas, colon, and lung, suggesting that oncogenic signaling is filtered through distinct tissue lineage-specific lenses shaped by developmental transcriptional states^10–12^. We hypothesize that the KRAS-driven oncogenic program is not a monolithic entity but is instead hardwired to the tissue-specific transcriptome defined by the cell of origin for a given tumor. Here, we describe the development of endogenous isogenic models across three permissive lineages and one non-permissive lineage to map the landscape of KRAS synthetic lethality. By integrating functional genomics with comparative transcriptome analyses, we demonstrate that oncogenic KRAS dependencies are profoundly divergent, underscoring the need for lineage-aware drug development in precision oncology.

## RESULTS

### Generation of Lineage-Specific Isogenic KRAS Mutant Models

To investigate the tissue-specific requirements for - mediated transformation, we engineered conditional, endogenous *KRAS* mutant isogenic lines in non-transformed, immortalized human cell lines. We selected models derived from “permissive” lineages, such as pancreas (HPNE^13^), colon (HCEC^14^), and lung (AALE^15^),as well as a “non-permissive” breast lineage (HMEC^16^) to serve as a biological contrast (**Fig. 1A**). We utilized a precise genome-editing pipeline to introduce a lox-Stop-lox (LSL) cassette^17^ into the endogenous *KRAS* locus, encoding KRAS G12C (lung) or KRAS G12D (colon, pancreas, and breast). Each mutant allele was fused with an N-terminal 3xHA tag to facilitate downstream detection (**Fig. 1B**).

**Figure 1.**
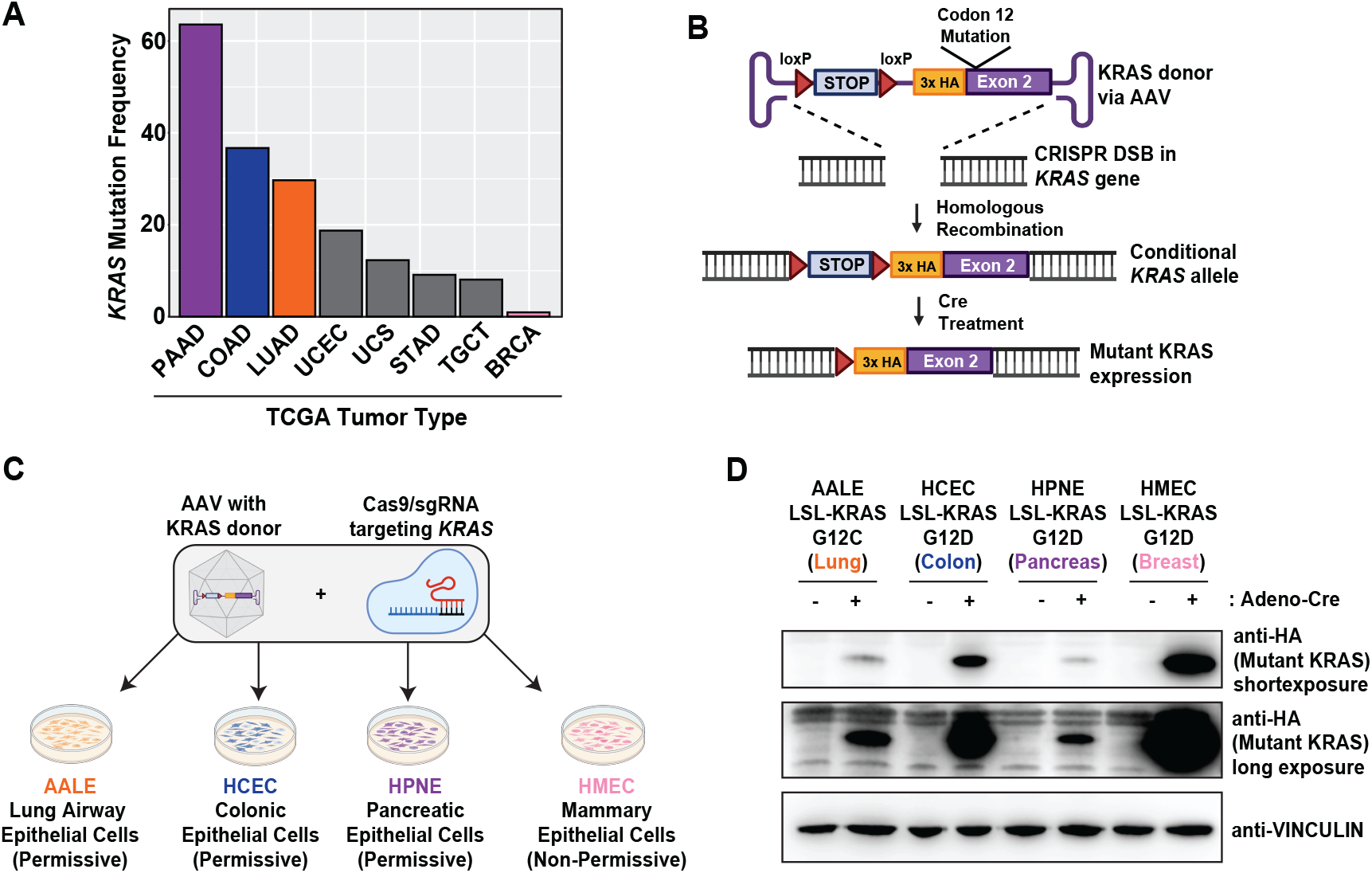
Isogenic KRAS cell line generation in permissive and non-permissive tissue lineages. **A**. *KRAS* mutational frequencies across TCGA tumor types. Bar chart representing the frequency of *KRAS* mutations across human cancers including Pancreatic Adenocarcinoma (PAAD), Colon Adenocarcinoma (COAD), Lung Adenocarcinoma (LUAD), Uterine Corpus Endometrial Carcinoma (UCEC), Uterine Carcinosarcoma (UCS), Stomach Adenocarcinoma (STAD), Testicular Germ Cell Tumors (TGCT), and Breast Invasive Carcinoma (BRCA). **B**. Schematic of the Adeno-associated virus (AAV) vectors cloned to include DNA homology surrounding a CRISPR/Cas9 induced double-strand break (DSB) near the start codon in exon 2 of the human *KRAS* gene. The construct features a loxP-STOP-loxP (LSL) cassette providing a transcriptional stop that is excised upon Cre recombinase delivery, permitting the endogenous expression of an N-terminal 3x-HA tagged mutant *KRAS*. **C**. Experimental workflow for engineering immortalized, non-transformed cell lines (AALE, HCEC, HPNE, and HMEC). Cells were nucleofected with a ribonucleoprotein (RNP) complex consisting of spCas9 and a *KRAS*-targeting sgRNA, followed by AAV-mediated delivery of the homology donor 24h post-nucleofection. **D**. Validation of inducible oncogene expression. Isogenic cells were transduced with adenovirus expressing either mCherry (control) or Cre recombinase. Whole-cell lysates were analyzed via Western blot using an anti-HA antibody to detect mutant *KRAS* protein, with VINCULIN serving as a loading control.

The engineering strategy involved nucleofection of recombinant spCas9 protein and sgRNAs targeting *KRAS* exon 2, followed by transduction with an adeno-associated virus (AAV) donor template to provide the repair construct for homologous recombination (**Fig. 1C**). Following puromycin selection and FLP-mediated excision of the selection cassette, single-cell clones were isolated and validated by genotyping PCR and Sanger sequencing to confirm the successful integration of the knock-in *KRAS* mutant allele and the preservation of the remaining *KRAS* WT allele.

To activate mutant KRAS expression, clones were treated with adenovirus expressing Cre recombinase, while adenovirus expressing mCherry served as a negative control. Western blot analysis confirmed the robust, inducible expression of the 3xHA-tagged mutant KRAS protein exclusively in Cre-treated cells (**Fig. 1D**). Of interest was the differing levels of KRAS produced in the different lines despite the identical chromosomal context. This isogenic system provides a controlled platform to isolate KRAS-specific genetic dependencies from the confounding genetic heterogeneity typically found in established cancer cell lines.

### In vitro characterization of the different KRAS Mutant Models

To determine how the lineage-specific context modulates oncogenic output, we evaluated downstream KRAS signaling across our isogenic pairs. Cells were transduced with either adeno-mCherry (control) or adeno-Cre to activate endogenous mutant KRAS, and activation of the canonical MAPK and PI3K/AKT pathways was assessed via western blot (**Fig. 2A**).

**Figure 2.**
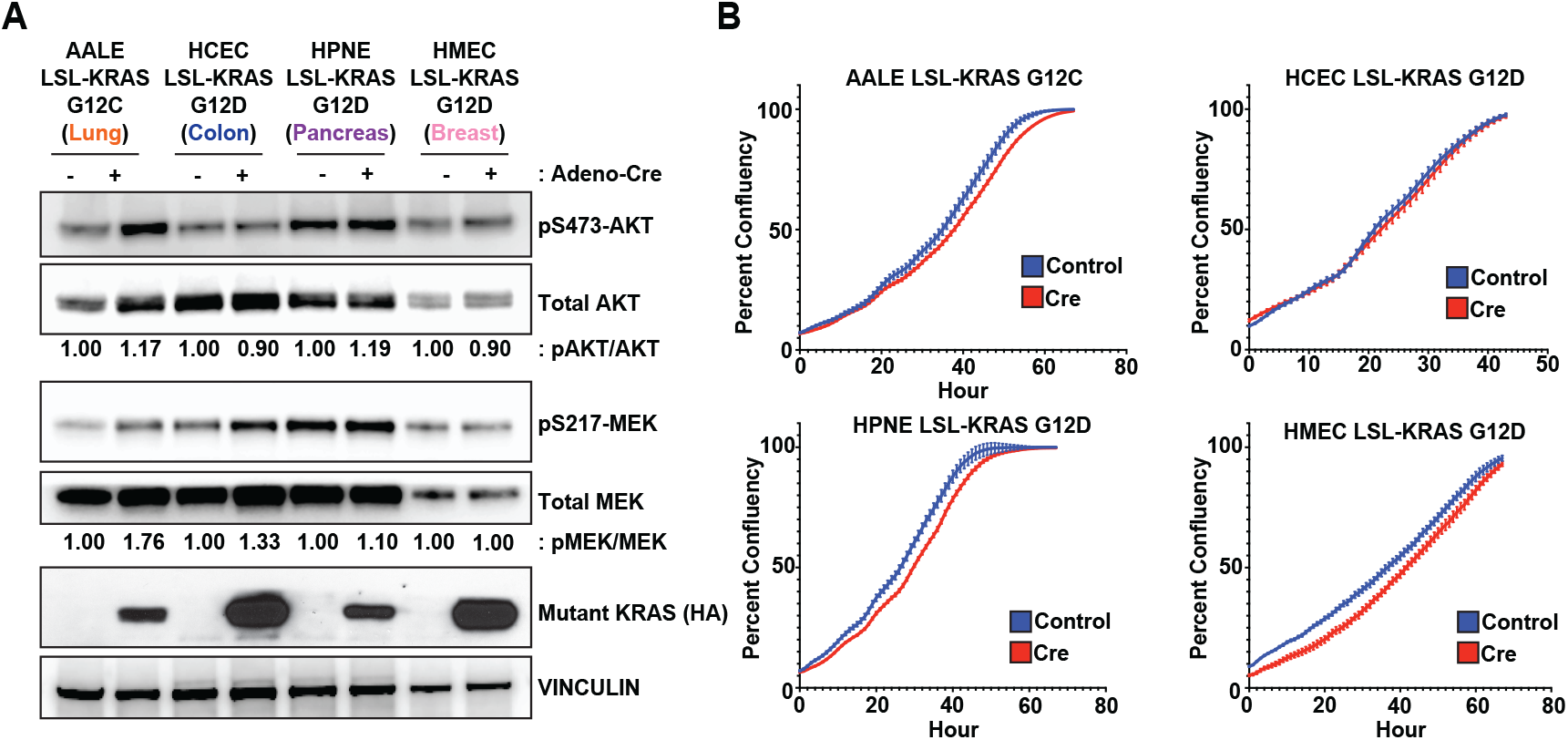
In vitro characterization of the isogenic KRAS cell lines. **A**. Downstream signaling upon mutant *KRAS* expression. Western blots of cell lysates from the four isogenic cell lines (AALE, HCEC, HPNE, and HMEC) treated with either mCherry control or Cre-expressing adenoviruses. Activation of the MAPK cascade (pSer217 MEK) and the PI3K/AKT pathway (pSer473 AKT) was evaluated to determine KRAS effector signaling upon endogenous KRAS mutant expression across different tissue lineages. Anti-HA was used to determine mutant KRAS expression. Total protein levels of MEK, AKT, and VINCULIN were used as loading controls. Numbers represent either phospho-AKT/total AKT or phospho-MEK/total MEK normalized to adeno-mCherry treatment within each cell lineage. **B**. Effects of KRAS mutant expression on cell proliferation. Isogenic cells treated with either mCherry control or Cre expressing adenoviruses were analyzed by live cell imaging to determine changes in cell proliferation over a time course of two to three days. Data points represent the mean with error bars for the SEM from three biological replicates.

In the permissive AALE and HCEC lineages, KRAS activation led to an appreciable increase in MAPK signaling, as evidenced by elevated levels of phospho-MEK (**Fig. 2A**). In the permissive AALE and HPNE cell line, activation of KRAS led to increased PI3K signaling as shown by an increase in phospho-AKT (**Fig. 2A**). In contrast, the non-permissive HMECs exhibited no detectable change in either MAPK or PI3K pathway activation following mutant KRAS expression (**Fig. 2A**).

Notably, the level of mutant KRAS protein, as measured by anti-HA immunoblotting, did not match the degree of effector pathway activation. Despite having the highest total expression of mutant KRAS, the non-permissive HMECs failed to activate the MAPK cascade (**Fig. 2A**). This disconnect between oncogene expression and signaling output suggests that lineage-specific negative feedback loops or the lack of necessary cofactors may suppress signaling in non-permissive contexts.

We next examined whether these biochemical changes translated into altered growth kinetics. Cell proliferation was monitored in real-time using Incucyte live-cell imaging to quantify confluency over time. Unexpectedly, the activation of mutant KRAS failed to significantly increase the proliferation rate of either the permissive or non-permissive cell lines under standard culture conditions (**Fig. 2B**). This observation is consistent with *KRAS* being a “primary hit” that requires additional genetic or microenvironmental context to achieve full morphological transformation and hyperproliferation in non-transformed, diploid human cells.

### Tissue Lineage-Specific Changes in the Transcriptome After KRAS Mutant Expression

To identify how KRAS mutant expression affects gene expression in permissive and non-permissive tissue lineages, we conducted RNA sequencing (RNA-seq) on each isogenic pair after inducing mutant KRAS. As might be expected, principal component analysis (PCA) showed that the main source of transcriptional variation was the tissue of origin rather than *KRAS* mutational status, with HCEC, AALE, HPNE, and HMEC lines grouping into distinct, lineage-specific clusters (**Fig. 3A**). We then examined each isogenic pair to compare WT and mutant KRAS transcriptomes within each tissue lineage (**Table S1**).

**Figure 3.**
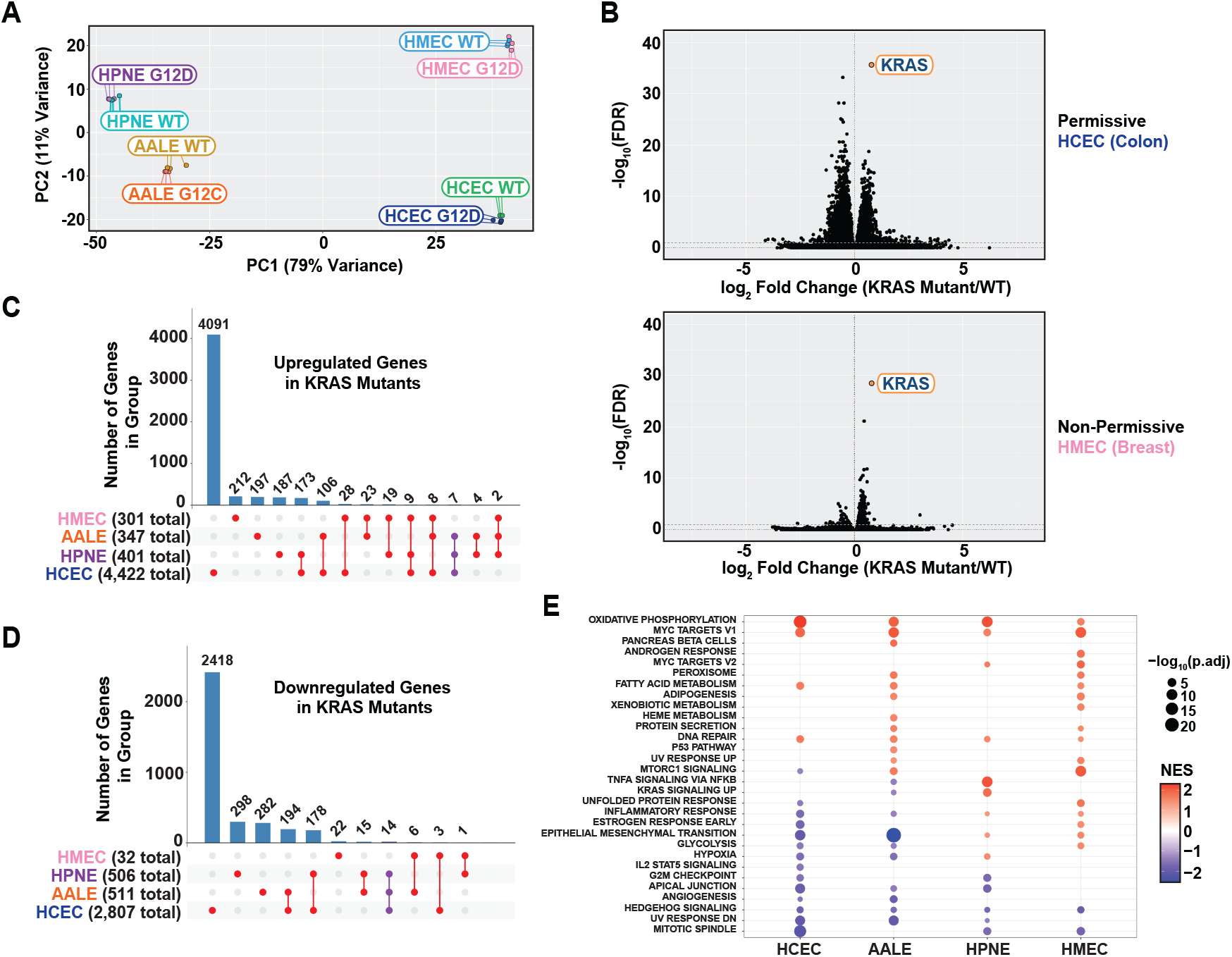
Transcription changes upon KRAS mutation induction in permissive and non-permissive tissue lineages. **A**. Principal component analysis (PCA) of RNA-seq transcriptomes from each isogenic pair (AALE, HCEC, HPNE, and HMEC) following treatment with either mCherry control (WT) or Cre (G12C/D) expressing adenoviruses. Samples cluster primarily by tissue of origin rather than mutational status, indicating that lineage identity remains the dominant driver of the global transcriptome. **B**. Volcano plots illustrating the magnitude and significance of differentially expressed genes (DEGs) following *KRAS* induction. The permissive HCEC line (top) exhibits massive transcriptional remodeling, whereas the non-permissive HMEC line (bottom) shows a markedly constrained response despite similar levels of oncogenic KRAS expression. **C**. Comparison of genes upregulated upon *KRAS* mutant expression. UpSet plot quantifying the overlap of significantly upregulated genes across all four isogenic pairs. Note the lack of a “universal” upregulated signature and the specific enrichment of only 7 genes shared among the three permissive lineages. **D**. Comparison of genes downregulated upon *KRAS* mutant expression. UpSet plot illustrating the overlap of significantly downregulated genes across each isogenic pair. No single gene was found to be downregulated in common across all four lines, with only 14 genes shared exclusively between the permissive lineages. **E**. Gene set enrichment analysis of Hallmark pathways upregulated or downregulated upon *KRAS* mutant expression. Dot plot showing some of the significantly enriched Hallmark pathways. Note the conservation of MYC target signatures and metabolic reprogramming (e.g. Oxidative Phosphorylation) across both permissive and non-permissive contexts and the lack of significantly downregulated pathways in the non-permissive HMECs.

Analysis of the pancreas derived HPNE cells revealed two-fold higher expression of both DUSP4 and DUSP6 in response to KRAS activation (**Table S1**), dual specificity kinases that are major negative regulators MAPK pathway activity. This could in part explain the lack of MAPK pathway activity observed in the HPNE cells upon KRAS activation (**Fig. 2A)**. The transcriptional recruitment of negative feedback regulators like DUSPs represents a potential lineage-specific ‘buffering capacity’ that can dictate a signaling threshold for KRAS-mediated transformation. Similarly, in non-permissive tissues like the HMECs, the absence of required lineage-defined co-factors may dampen signaling, leading to reduced MAPK activation despite robust mutant KRAS expression. For example, the levels of AKT and MEK are significantly lower in HMEC cells compared to AALE, HCEC and HPNE (Figure 2A).

The colonic epithelial HCEC cells exhibited a massive transcriptional response, with 4,422 significantly upregulated genes, a figure nearly fifteen-fold higher than the other permissive lines (347 for AALE, 401 for HPNE) and the non-permissive HMECs (301) (**Fig. 3C**). Interestingly, while mutant KRAS expression was highest in HCEC cells, comparable only to the non-permissive HMEC line (**Fig. 1D**), the magnitude of transcriptional output did not scale linearly with oncogene levels across lineages. In both HCEC and HMEC, KRAS was the top significantly upregulated gene (**Fig. 3B**); however, the HCEC permissive context appears uniquely poised to amplify this signal into a broad transcriptomic shift. Despite reduced levels of mutant KRAS expression (**Fig. 2A**), the permissive AALE and HPNE cell lines also responded more robustly than the non-permissive HMEC lineage (**Fig. S1**). These data further underscore that “permissivity” is not merely the presence of the mutation, but the innate ability of the tissue-specific regulatory landscape to propagate oncogenic signaling into a diverse downstream program.

In addition to the induction of oncogenic programs, we analyzed the landscape of transcriptional repression following KRAS activation. Consistent with the trends observed in upregulated transcripts, the colonic epithelial HCEC line exhibited the most profound shifts, with 2,807 significantly downregulated genes (**Fig. 3D**). This vast suppression of the transcriptome in HCECs stands in stark contrast to the more tempered responses in the lung (AALE, 511 genes) and pancreas (HPNE, 506 genes) (**Fig. 3D**). Most notably, the non-permissive HMEC line was remarkably resistant to KRAS-mediated gene repression, with only 32 genes significantly downregulated (**Fig. 3D**).

Analysis of upregulated transcripts after oncogenic KRAS expression revealed no common genes shared between each of our four isogenic cell line pairs (**Fig. 3C**). In fact, only seven genes were shared between our three permissive lineages (*AL450405*.*1, RBBP9, RPL36A-HNRNPH2, ARHGDIB, RPL36A, S100A10, POLD2*). Four of these upregulated transcripts are involved in protein translation (AL450405.1, RBBP9, RPL36A-HNRNPH2, and RPL36A), indicating a ‘hard-wired’ biosynthetic requirement for KRAS-mediated transformation that transcends tissue boundaries (**Table S1**).

A comparative analysis across all four lineages revealed no single downregulated gene in common (**Fig. 3D**), further reinforcing the hypothesis that mutant KRAS does not dictate a universal repressive transcriptional program. Even when restricting the analysis to the three permissive lineages, we identified only 14 shared downregulated genes (**Fig. 3D**). This small subset of common downregulated factors includes several genes involved in cell-cell adhesion, extracellular matrix (ECM) organization, and growth factor regulation, such as *FZD2, LRIG3, PCDH18*, and *DARS2*.

The disproportionate scale of gene repression in HCEC cells, nearly 90 times that of non-permissive HMECs, suggests that in permissive contexts, KRAS activation triggers a lineage-specific remodeling of the cellular environment. The suppression of genes like LRIG3, a known negative regulator of receptor tyrosine kinase (RTK) signaling, and PCDH18, a protocadherin associated with tissue architecture, may represent a coordinated effort to remove biological “brakes” that otherwise prevent transformation. The failure of KRAS to achieve this degree of transcriptional remodeling in HMECs likely contributes to the non-permissive nature of the breast lineage, where the original cellular identity remains largely intact and resistant to oncogenic reprogramming by mutant KRAS.

Gene set enrichment analysis (GSEA) of pathways upregulated in KRAS mutant cells revealed increased expression of genes related to key metabolic pathways, notably for protein production (e.g. *the ribosome* and *protein secretion*) and major metabolic pathways (e.g. *oxidative phosphorylation* and *fatty acid metabolism*) (**Fig. 3E and Table S1**). These gene sets are largely composed of genes upregulated with increased MYC transcriptional activity. Not surprisingly, the “MYC targets V1” gene set was among the highest-scoring pathways upregulated by mutant KRAS across all 4 cell models. These data suggest that MYC transcriptional activity is a crucial signaling node downstream of mutant KRAS, regardless of tissue lineage.

Intriguingly, GSEA also revealed a conservation of downregulated pathways, specifically those governing the cell cycle and mitosis (e.g., *Mitotic Spindle, G2M Checkpoint*, and *E2F Targets*) (**Fig. 3E**). While oncogenic KRAS is typically associated with accelerated growth, the suppression of these hallmarks suggests an immediate activation of compensatory growth-arrest programs. Because our models utilize non-transformed, telomerase-immortalized cells rather than established cancer cell lines, they likely retain intact tumor-suppressive feedback loops. This phenomenon is reminiscent of Oncogene-Induced Senescence (OIS), in which the sudden hyperactivation of the MAPK pathway triggers a cell cycle exit to prevent the propagation of potential malignant cells. The strong negative enrichment of E2F targets, a hallmark of the G1/S transition, suggests that these cells may be activating a transcriptional response to counteract KRAS-driven overproliferation.

### Identification of Tissue Lineage-Specific Synthetic Lethal Interactions

To map the genetic dependencies of oncogenic *KRAS* across different lineages, we performed genome-wide CRISPR/Cas9 screens in KRAS WT (mCherry) and KRAS mutant (Cre) isogenic pairs for each permissive tissue lineage (**Fig. 4A**) and the non-permissive HMEC cells for comparison. Following lentiviral transduction of a pooled gRNA library and puromycin selection, cells were harvested at population doubling 0 (PD0) and PD10. We quantified sgRNA abundance using next-generation sequencing (NGS) to identify synthetic lethal (SL) candidates, i.e. genes whose depletion selectively impaired the fitness of *KRAS* mutant cells relative to their *KRAS* WT counterparts.

**Figure 4.**
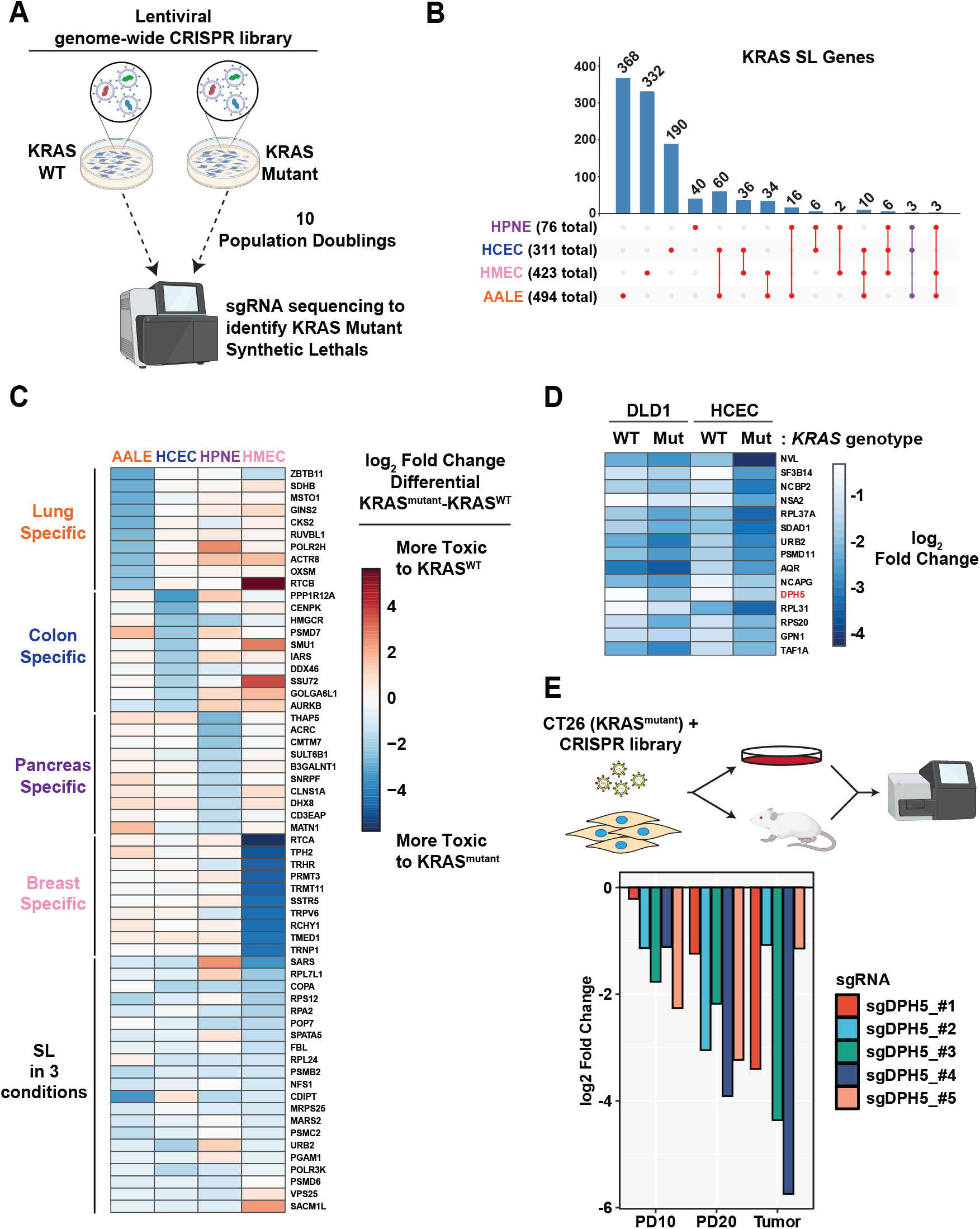
Synthetic lethal interactions with oncogenic KRAS are remarkably tissue specific. **A**. Schematic outlining our genome-wide CRISPR screens in each isogenic pair. A pooled lentiviral library targeting ∼18,200 human genes (5 sgRNAs per gene) was transduced into mCherry control (KRAS WT) and Cre-expressing (KRAS Mutant) adenovirus-treated cells. Following selection, an initial reference population was harvested at population doubling zero (PD0). Cells were maintained for an additional 10 PDs, after which genomic DNA was harvested and sgRNA abundance was quantified via next-generation sequencing to identify KRAS synthetic lethal (SL) hits. **B**. UpSet plot showing the overlaps of KRAS SL genes from each of the isogenic pairs. Comparisons of significantly scoring KRAS SL hits across lung, colon, pancreas, and breast lineages. The diagram highlights the minimal overlap (only 3 shared genes, connected by the purple line) among permissive lineages, illustrating the profound influence of the tissue-specific landscape on oncogenic KRAS dependency. **C**. Heatmap demonstrating the top tissue lineage specific KRAS SL genes and the KRAS SL genes that scored in 3 different cell lines. **D**. Comparison of KRAS SL genes in DLD1 human colorectal cancer (CRC) cells and the HCEC isogenic pair. Heatmap demonstrating the top 15 KRAS SL genes identified in both the DLD1 and HCEC cell lines including the diphthamide pathway gene *DPH5*. **E**. DPH5 is required for KRAS mutant mouse CRC cell growth in vitro and in vivo. Analysis of CRISPR screening data in CT26 mouse tumor cells grown in vitro or subcutaneously in BALB/c mice. Bar graph of five different *Dph5*-targeting sgRNAs demonstrating the requirement of DPH5 expression for both in vitro and in vivo tumor growth.

Our analysis revealed a striking divergence in *KRAS* dependencies. We identified 76 SL genes in pancreatic cells, 494 in lung cells, 311 in colorectal cells, and 423 in the breast cells (**Fig. 4B**). Remarkably, only three genes (*PSMD6, VPS25*, and *SACM1L*) were shared across all three permissive tissue types. The overlap between individual lineages was unexpectedly low, ranging from only 4.5% to 29.7%, underscoring the profound influence of tissue context on *KRAS* mutant vulnerability (**Fig. 4B, Table S2**). Comparison of HMEC dependencies with the permissive lineages revealed minimal overlap (47 with lung, 52 with colon, and 11 with pancreas). Based on our KRAS SL score, we identified the top candidate tissue-lineage-specific KRAS SL genes (**Fig. 4C**). Comparison of KRAS SL hits between tissue types using GSEA identified that mutant lung cells exhibit an elevated sensitivity to mitochondrial translation inhibition (**Table S2**), consistent with our previous report^9^. While a small subset of common hits converged on core homeostatic processes, including mRNA processing, cell division, and translation, most SL interactions were lineage-restricted.

We next analyzed our previous KRAS SL CRISPR data in isogenic pairs of DLD1 colorectal cancer (CRC) cells. We identified candidate KRAS SL hits that were in common between the DLD1 and HCEC isogenic pairs including the diphthamide synthesis gene *DPH5* (**Fig. 4D**). We analyzed our previous in vivo tumor screens performed using *KRAS* mutant CT26 murine CRC cells and identified *Dph5* as a requirement for CT26 proliferation and tumorigenesis across 5 independent sgRNAs (**Fig. 4E**). Further analysis of our CRISPR screening results revealed diphthamide synthesis genes are KRAS SL hits in our other isogenic cells including *DPH2* and *DPH3* in the AALE, *DPH5* and *DPH6* in HCEC, and *DPH2* in the HMEC (**Table S2**). We analyzed results from the Cancer Dependency Map (DepMap), a collection of whole-genome CRISPR screens across human tumor cell lines including 237 cell lines derived from lung, colon, and pancreas. We separated gene dependency probability scores based on *KRAS* mutation status for these cell lines to identify the genes whose knockout is synthetically lethal in *KRAS* mutant cells (**Table S3**). Not surprisingly, the top scoring KRAS mutant dependency is KRAS itself (**Fig. S2A**). Potential *KRAS* mutant SL genes include Ras signaling pathway components like c-Raf (*RAF1*) along with previously described KRAS SL genes including *EFR3A*^18^ and *DOCK5*^8^ (**Fig. S2B**). Analysis of Chronos gene essentiality scores in each cell line showed that *KRAS* and *RAF1* knockout is more toxic in KRAS mutant cells as expected (**Fig. S2C-D**). Analysis of essentiality scores for DPH5 and the diphthamide pathway revealed that KRAS mutant cells are more sensitive to their loss (**Fig. S2E-F**) supporting our isogenic screening results in established human tumor cell lines. Collectively, these data from human isogenic models, established cancer cell lines, murine tumor models, and large-scale human tumor cell CRISPR screens underscore a requirement for diphthamide synthesis to sustain KRAS-driven oncogenic growth.

These findings suggest that *KRAS* mutant synthetic lethality is not a universal property of the oncogene itself but is instead hardwired into the transcriptional and epigenetic landscape of the cell of origin. Even when comparing permissive to non-permissive contexts, the genetic networks required to sustain oncogenic KRAS signaling remain fundamentally divergent, suggesting that therapeutic targets identified in one organ may fail to translate to others.

## DISCUSSION

The clinical success of PARP inhibitors in BRCA-mutant malignancies has solidified synthetic lethality as a cornerstone of precision oncology. However, applying this paradigm to KRAS mutations has proven difficult, largely due to the elusive nature of any “universal” dependencies. Our study demonstrates that KRAS synthetic lethality is not an intrinsic property of the oncogene itself, but rather an emergent property of the oncogene interacting with the specific proteome and transcriptome developmentally wired by the identity of the tissue lineage.

By utilizing non-transformed, isogenic models, we isolated the immediate consequences of KRAS activation. The surprising lack of overlap in SL hits, with only three genes shared across lung, colon, and pancreas, suggests that the “KRAS-addiction” state is wired through divergent pathways that are highly dependent on the cell of origin. For instance, the unique dependency of lung cells on mitochondrial translation may reflect the specific metabolic or oxidative stress thresholds inherent to pulmonary epithelium, which are absent in the ductal environment of the pancreas.

Our gene-expression analyses revealed a common MYC-driven signature across our isogenic cell lines that remodels the transcriptome upon KRAS activation. While the induction of these pathways appears universal and sustained MYC signaling downstream of mutant KRAS is required for tumorigenesis, the specific genetic dependencies required to maintain this metabolic state remain highly context-dependent, as evidenced by our genomic screens. Nevertheless, the induction of MYC-driven metabolic pathways creates a targetable translational stress node that can be exploited to treat KRAS mutant cells. Previous studies have demonstrated that inhibition of protein translation pathways are potent blockers of KRAS-driven tumor growth in different tumor types. Our data further demonstrate that targeting specific translational machinery, like the mitochondrial ribosome in lung cells or the diphthamide pathway in colon cells. leads to robust growth inhibition in the context of a *KRAS* mutation.

Our identification of the diphthamide synthesis pathway as a *KRAS* mutant dependency provides a mechanistic anchor for this lineage-restricted model. Diphthamide is a unique post-translational modification of eukaryotic elongation factor 2 (eEF2) that is essential for ribosomal translocation and translational fidelity. Notably, the requirement for this pathway in RAS-driven growth appears evolutionarily conserved; previous studies demonstrated that Ras-induced hyperplasia in the *Drosophila* gut requires a functional diphthamide synthesis pathway^19^ and oncogenic RAS signaling has been shown to modulate cellular sensitivity to diphtheria toxin, which targets the diphthamide residue^20^. Our data reveal that *KRAS* activation, particularly in the colonic lineage (HCECs), triggers a massive transcriptional amplification of this biosynthetic machinery. This likely creates a ‘hyper-translational’ state that, while supporting oncogenic growth, simultaneously imposes a rigid dependency on *DPH5*.

By integrating our transcriptomic and CRISPR screening datasets, we propose that the KRAS-driven ‘permissivity’ creates a biosynthetic bottleneck. While our non-transformed cell models suppress core mitotic programs likely as a defense mechanism (**Fig. 3E**), they simultaneously become more dependent on the translation machinery, such as the diphthamide synthesis pathway (Fig. 4D), to navigate the resulting replication stress and sustain cell growth.

In the same way that cell proliferation shows profound tissue specificity upon genetic perturbation in gain-of-function screens^21^, synthetic lethality shows an equivalent magnitude of tissue specificity. This has ramifications for therapeutics that target synthetic lethal partners rather than the oncogene itself. Additionally, this work suggests that basket trials, which enroll patients based solely on *KRAS* mutational status, may be problematic due to tissue specificity issues. Thus, future drug development will need to consider “lineage-aware” combinations to achieve clinical efficacy.

## MATERIALS AND METHODS

### Cell Culture

Human colonic epithelial cells (HCECs) were obtained from Dr. Jerry Shay (UT-Southwestern) and have been described previously^14^. Human airway epithelial cells (AALE) were obtained from Dr. Kevin Haigis (DFCI) and have been described previously^15^. Human nestin-positive pancreatic epithelial cells (HPNEs) were obtained from Dr. Channing Der (UNC) and have been described previously^13^. Human mammary epithelial cells (HMECs) were from a reduction mammoplasty and have been described previously^16^. HCEC cells were grown in a mixture of DMEM (Thermo cat# 10569010) and Medium 199 (Sigma cat# M4530) at a 4:1 DMEM:Medium 199 ratio supplemented with 2% fetal calf serum (FCS) (Corning), 20 ng/mL EGF (Thermo cat# PHG0311), 1 μg/mL hydrocortisone (Sigma cat# H4001), 10 μg/mL insulin (Sigma cat# I0516), 2 μg/mL transferrin (Sigma cat# T8158), and 5 nM sodium selenite (Sigma cat# S5261), and 1% penicillin/streptomycin (Pen/Strep Thermo cat# 15140122). AALE cells were grown in SABM basal media (Lonza cat# CC-3119) supplemented with SAGM singlequots supplements (Lonza cat# CC-4124). HPNE cells were grown in a mixture of DMEM and M3:BaseF (InCell cat #M300F) medias at a ratio of 3:1 DMEM:M3:BaseF supplemented with 10% FCS and 1% Pen/Strep. HMEC cells were grown in MEGM basal media (Lonza cat# CC-3151) supplemented with MEGM SingleQuots Supplement (Lonza cat# CC-4136). HEK293T cells were used to generate adeno-associated virus (AAV) and lentiviruses and were maintained in DMEM supplemented with 10% FCS and 1% Pen/Strep. All cells were maintained at 37°C and 5% CO_2_.

### Adeno-associated Virus Production

Recombinant AAV serotype 2 was packaged in 293T cells using the pHelper and pRC2-mi342 plasmids (Takara cat# 6652). A 10-cm plate of 293T cells was transfected with 5 μg pHelper, 5 μg pRC-mi342, and 5 μg LSL-3xHA KRAS targeting vector plus 45 μL PolyJet transfection reagent (SignaGen cat# SL100688). Three days after transfection, 293T cells were harvested by scraping and AAV2 was isolated using AAVpro Extraction Solution (Takara cat# 6235). AAV was frozen at −80°C until use.

### LSL-3xHA KRAS Knock-in Cell Line Generation

One million AALE, HCEC, and HPNE cells were transfected in a Lonza 4D nucleofector (Lonza cat# AAF-1003X) using a 1 mL cuvette with 82 μL of SE solution and 18 μL Supplement 1 (Lonza cat #V4XC-1024) 1.5 μg of pcDNA3.3 TOPO spCas9 (Addgene cat# 41815) and 500 ng of a U6-driven KRAS sgRNA plasmid (5′-TGACGATACAGCTAATTCA-3′) to target exon 2 of human *KRAS*. Pulse codes EN-113, DN-100, and CM-130 were used for HCEC, HPNE, and AALE cells, respectively. Cells were plated in a 6-well plate post-nucleofection and allowed to recover for 16 h. After recovery, 100 μL of AAV LSL-3xHA KRAS targeting donor was added and incubated on cells overnight. Five days after AAV delivery, successfully infected cells were selected with 1 μg/mL puromycin (Thermo cat# A1113803) until a negative control non-infected plate was completely killed. After puromycin selection, cells were infected with Adeno-FLPe recombinase (Vector Biolabs cat# 1760) to excise the puromycin selection cassette and successfully recombined cells were isolated with 10 μM ganciclovir selection. Single cell clones were then screened by Adeno-Cre (Vector Biolabs cat# 1773) treatment to identify clones that had successfully integrated the LSL cassette, 3xHA tag onto the amino terminus of endogenous KRAS, and either the G12C or G12D mutation. All clones were verified by PCR from genomic DNA and Sanger sequencing of the *KRAS* knock-in allele. The non-targeted allele was verified to be non-mutated by Sanger sequencing of PCR amplicons. Plasmid maps for the KRAS sgRNA and AAV targeting vectors can be found in the Reagents section.

### Western Blots

All cells were lysed in 1X RIPA lysis buffer (Boston Bioproducts cat# BP-115X) with protease and phosphatase inhibitors (Thermo cat# 78445). Protein was quantified by Bradford assay (Bio-Rad cat# 5000006). Protein lysates were separated on 4-12% Bis-Tris gels (Thermo cat# NP0322BOX) and transferred to nitrocellulose membranes (Bio-Rad cat# 1704158) using a Bio-Rad Transblot transfer device. Membranes were blocked with 5% milk (Fisher cat# NC9121673) in TBST (Cell Signaling cat# 9997S) and incubated with the following antibodies for 1h at room temperature or overnight at 4°C: anti-HA HRP (Cell Signaling cat# 2999S) and anti-Vinculin (Sigma cat# V9131). For anti-Vinculin, a secondary antibody (Jackson ImmunoResearch, goat anti-mouse cat# 115-035-003) was used at 1:5000 for 1 h at room temperature. Blots were washed three times with TBST following secondary antibody incubation, developed with ECL (Revvity cat# NEL103001EA), and visualized on a LiCOR Odyssey imager.

### CRISPR Screens

A library of gRNAs targeting 18,166 genes with five gRNAs per gene for a total of ∼91,000 gRNAs was synthesized and cloned into the lentiCRISPR V2 puro vector. Pooled virus was prepared by transfecting 293T cells with the library plasmid pool with psPax2 and pMD2.G lentiviral packaging vectors (Addgene cat# 12260 and 12259). Viral supernatants were harvested at 48 and 72 h post-transfection and concentrated with lenti-X concentrator solution (Takara cat # 631231). AALE, HCEC, and HPNE cells were infected at a low MOI (0.2) with a representation of 500 in triplicate. Cells were selected with puromycin (1 or 2 μg/mL) for 3 d until an uninfected control plate completely died. An initial cell pellet was taken as PD0, cells were grown an additional PD10, and a final cell pellet was collected. Genomic DNA was isolated by phenol/chloroform (Thermo cat# 15593031) extraction and gRNAs were PCR amplified with barcoded primers for sequencing on an Illumina NextSeq 500. Sequencing reads were aligned to the initial library and counts were obtained for each gRNA (**Table S2**).

### RNA-sequencing

Cell cultures grown to ∼60% confluency were collected in triplicate for each isogenic cell line pair, either treated with control adenovirus or Cre recombinase to initiate mutant K-Ras expression. Cells were harvested by scraping on ice in PBS and mRNA was isolated using a RNeasy mRNA extraction kit (Qiagen). Isolated mRNA was sent out for library prep and RNA-sequencing (BGI). Sequencing data were downloaded and aligned to the human reference transcriptome (GRCh38) to generate a read-count table for downstream differential expression analysis. Raw read counts and analyses can be found in **Table S1**.

### Statistical Analyses

MAGeCK^22^ and edgeR^23^ were used to calculate p values, FDRs, and log_2_ fold changes (log_2_FC) for comparison between the PD10 and PD0 samples for each cell line. The log_2_FC for each gene was determined by the mean log_2_ fold change (PD10/PD0) of the five sgRNAs targeting that gene. MAGeCK analyzed data were used to identify *KRAS* mutant synthetic lethal genes as defined by a differential log_2_ fold change (log_2_FC *KRAS* mutant – log_2_FC *KRAS* WT) of at least −0.5 and where the gene in the *KRAS* mutant cell line had an adjusted p-value (FDR) < 0.2 for depletion of that gene (**Table S1**). GSEA analysis was performed using DAVID (https://davidbioinformatics.nih.gov/home.jsp). For RNA-seq analyses, raw read counts were collapsed to the gene level and analyzed using DESeq2^24^.

### Cancer DepMap Analysis

Whole-genome CRISPR screening data were obtained from the DepMap portal [Release 25Q3]. Gene dependency scores (Post-Chronos gene dependency probability estimates) were compared between KRAS WT and KRAS mutant (including both heterozygous and homozygous mutants) cell lines across 237 (120 KRAS WT and 117 KRAS mutant) lung, colon, and pancreatic cell lines. Differential dependency was assessed using a two-tailed Wilcoxon rank-sum test, and p-values were adjusted for multiple hypothesis testing using the Benjamini-Hochberg (BH) method.

## ACKNOWLEDGMENTS

This work was supported by a grant from the Cancer Research Grand Challenge and the Mark Foundation for Cancer Research to the SPECIFICANCER Team, The Ludwig Family Foundation. The authors wish to acknowledge Eric Wooten for computational guidance during screen analyses, Jerry Shay for providing the HCEC line, Kevin Haigis for providing the AALE line and helpful discussions about KRAS biology, and Channing Der for providing the HPNE line. S.J.E. is an Investigator for the Howard Hughes Medical Institute.

## Author Contributions

T.D.M, M.Y.C, and J.C.M performed all experiments. T.D.M and S.J.E designed the experiments, analyzed data, and wrote the manuscript with input from all authors.

---

**Figure S1.**
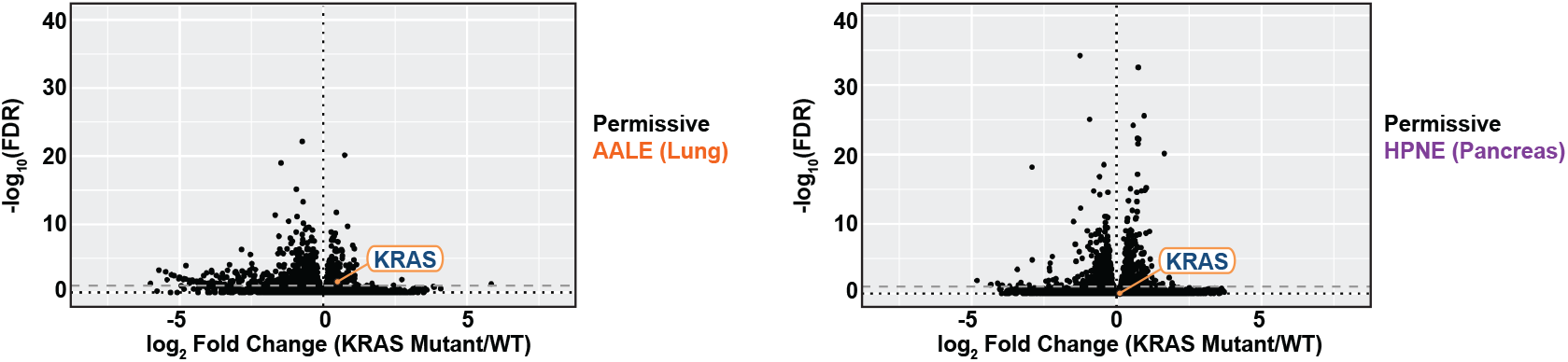
Transcriptome changes in permissive AALE and HPNE cell lines. **A**. Volcano plots illustrating the magnitude and significance of differentially expressed genes (DEGs) following *KRAS* induction in permissive AALE (left) and HPNE (right) cell lines.

**Figure S2.**
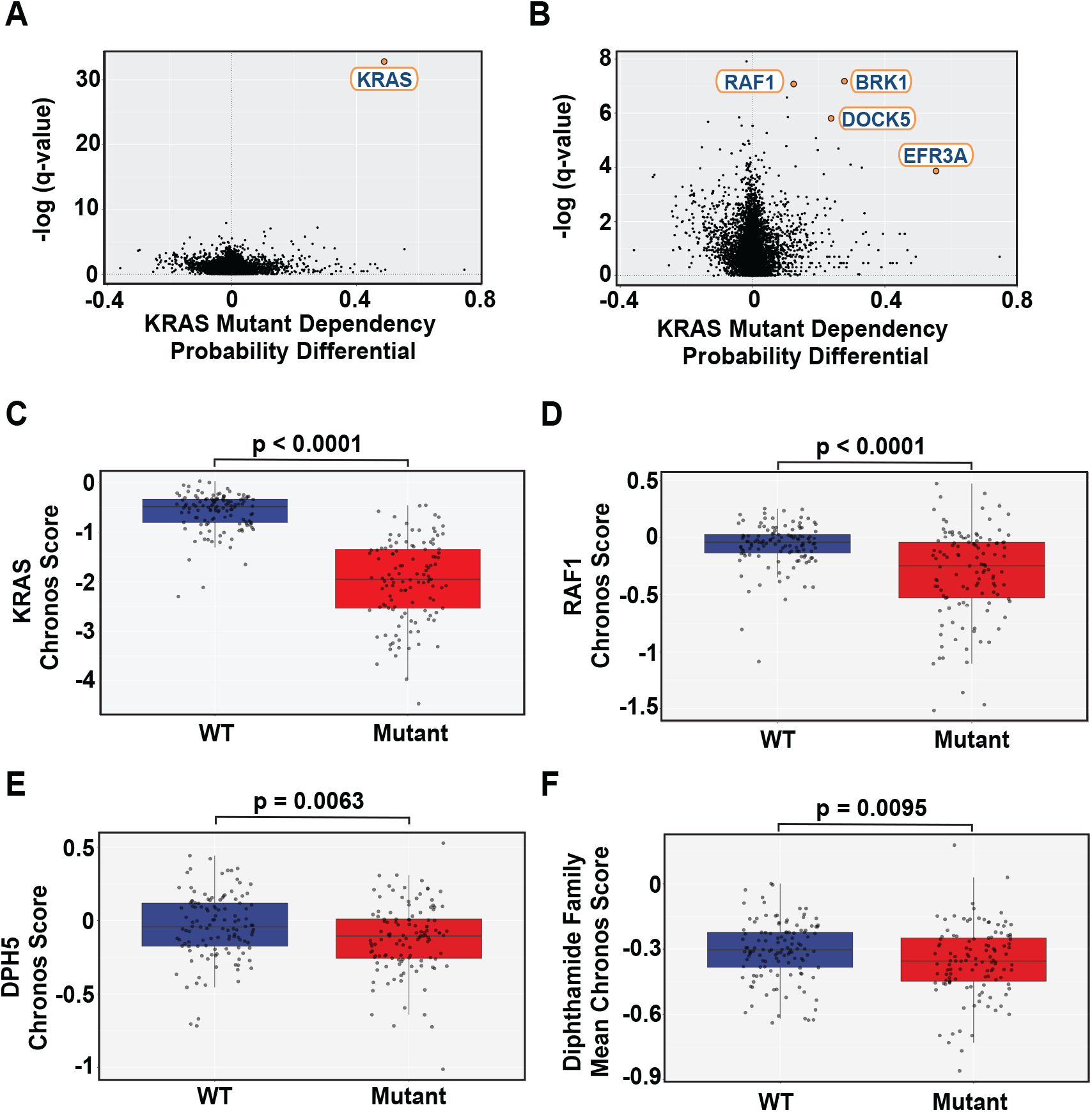
Analysis of KRAS mutant gene dependencies from the Cancer DepMap. **A**. Gene dependency differences in KRAS WT and mutant human colon, lung, and pancreatic tumor cells. Post-Chronos gene dependency probability estimates were compared between KRAS WT and KRAS mutant cell lines from whole-genome CRISPR screens. A differential of the probability estimate was determined by subtracting the KRAS WT probability from the KRAS mutant probability where values closer to +1 indicate more essential in KRAS mutant cells and −1 indicate more essential in KRAS WT cells. **B**. Gene dependency plot after removing the KRAS gene itself to provide a zoomed in view of KRAS mutant dependencies. **C**. Loss of KRAS is more toxic to KRAS mutant cell lines. **D**. Loss of c-Raf is more toxic in KRAS mutant cells. **E**. KRAS mutant cells are more sensitive to DPH5 knockout. **F**. KRAS mutant cells are more sensitive to inhibition of the diphthamide synthesis pathway.

## SUPPLEMENTAL FILES

**Table S1. RNA-sequencing data and analyses from the isogenic KRAS cell lines** Raw and normalized read counts for each gene, DESeq2 analyses comparing *KRAS* WT to *KRAS* mutant cells, upregulated and downregulated genes (FDR < 0.05), and GSEA for each cell line.

**Table S2**. CRISPR screening data and analyses from the isogenic KRAS cell lines Raw sgRNA read counts, MAGeCK and edgeR analyses for KRAS WT and KRAS mutant CRISPR screens, KRAS SL gene overlaps between tissue lineages, and GSEA of KRAS SL genes for each cell line.

**Table S3. Cancer Dependency Map (DepMap) analysis to identify putative KRAS SL genes across lung, colon, and pancreas cells**. Gene dependency scores for KRAS WT and KRAS mutant cells and adjusted p-values based on whole-genome CRISPR screens.

**Plasmid maps/sequences for the KRAS exon 2 targeting sgRNA and the AAV homology donors for LSL-3xHA KRAS G12C or G12D knock-in**.

